# A thousand metagenome-assembled genomes of *Akkermansia* reveal new phylogroups and geographical and functional variations in human gut

**DOI:** 10.1101/2020.09.10.292292

**Authors:** Qing-Bo Lv, Sheng-Hui Li, Yue Zhang, Yan-Chun Wang, Yong-Zheng Peng, Xiao-Xuan Zhang

**Affiliations:** College of Veterinary Medicine, Qingdao Agricultural University, Qingdao 266109, China; College of Animal Science & Veterinary Medicine, Heilongjiang Bayi Agricultural University, Daqing 163319, China; Shenzhen Puensum Genetech Institute, Shenzhen 518052, China; College of Animal Science and Technology, Jilin Agricultural University, Changchun 130118, China; Department of Laboratory Medicine, Zhujiang Hospital, Southern Medical University, Guangzhou 510282, China

**Keywords:** *Akkermansia muciniphila*, Metagenome-assembled genome, Population structure, Geographical variation, Functional specificity, Gut microbiota

## Abstract

The present study revealed the genomic architecture of *Akkermansia* in human gut by analyzing 1,119 near-complete metagenome-assembled genomes, 84 public available genomes, and 1 newly sequenced *A. glycaniphila* strain. We found that 1) the genomes of *Akkermansia* formed 4 species (including 2 candidate species) with distinct interspecies similarity and differed genomic characteristics, and 2) the population of *A. muciniphila* was structured by 3 previously identified phylogroups (Amuc I, II, and III) referring to 1,132 genomes and 1 new phylogroup (defined as Amuc IV) that contained 62 genomes. Amuc III was presented in Chinese population and Amuc IV was mainly distributed in western populations. A large number of gene of functions, pathways, and carbohydrate active enzymes that specifically associated to phylogroups. Our findings based on over a thousand genomes strengthened the previous knowledge and provided new insights into the population structure and ecology of *Akkermansia* in human gut.

## Introduction

*Akkermansia* is a well-focused genus that has been regarded as a representative of the phylum Verrucomicrobia in the human and animal gut [1, 2]. To date, only two species of *Akkermansia, A. muciniphila* and *A. glycaniphila* [2, 3], are isolated and comprehensive described. *A. muciniphila* is widely present in the intestinal mucosa of human [4-7], and it can degrade the highly glycosylated proteins in epithelial mucosa and produce diverse structural molecules such as short-chain fatty acids (SCFAs) [2]. The host range of *Akkermansia* genus is wide, ranging from mammals (mainly *A. muciniphila*) to non-mammals (e.g. *A. glycaniphila* is isolated from python [3]) that differed greatly in physiology, dietary structure and the composition of mucinous proteins in the gut [8]. There is growing evidence that *A. muciniphila* is an excellent candidate probiotic. Previous studies have shown the health-promoting effects of *A. muciniphila* [9, 10], as relative abundance of *A. muciniphila* in gut microbiota were negatively correlated with multiple metabolic disorders such as hyperlipidemia [11], severe obesity [12], and type 2 diabetes [13]. Furthermore, supplementation with *A. muciniphila* in mice exerted a protective effect on mice colitis induced by dextran sulfate sodium, and prevented the age-related decline in thickness of the colonic mucus layer [14, 15]. In clinical trials, oral supplementation *A. muciniphila* was considered as a safe and well tolerated intervention for weight losing, that improved insulin sensitivity and reduced insulinemia and plasma total cholesterol [16].

The genome research of *Akkermansia* is relatively lack, and many functional specificities of the *Akkermansia* genus remain unclear. After investigating a large number of full-length 16S sequences in 2011, it had been found that at least 8 species of *Akkermansia* genus resident in human digestive tract [17]. However, only two strains, *Akkermansia muciniphila* ATCC BAA-835 and *Akkermansia glycaniphila* Pyt [17, 18], were whole-genomic sequenced until 2017. As a great expand, in our previous study [19], we sequenced and analyzed the draft genomes of 39 *A. muciniphila* strains isolated from China and divided the population structure of this species into three phylogroups (Amuc I, Amuc II, and Amuc III). These phylogroups showed high genetic diversity presenting distinct metabolic and functional features. However, these strains were clearly not strong representation, giving the widespread global distribution of *A. muciniphila*, especially in the western population. In 2019, Pasolli *et. al*. [20] reconstructed over 150,000 microbial genomes via 9,428 metagenomes and expanded the pangenomes of human-associated microbes. These metagenome-assembled genomes (MAGs) included over one thousand high quality *Akkermansia* draft genomes, which provides the potential for studying the structure of *Akkermansia* across countries and populations. In this study, by analyzing these MAGs and combining publicly available genomes as well as a newly sequenced *Akkermansia* strain, we comprehensively revealed the population structure of the genus *Akkermansia* and discovered a new species-level *A. muciniphila* phylogroup (Amuc IV) and two candidate species (*Akkermansia* sp. V and *Akkermansia* sp. VI). We also investigated the functional specificity of *Akkermansia* species and phylogroups. Our study reinforces previous findings and provides new insights into the research on *Akkermansia*.

## Results and Discussion

### Metagenome-assembled genomes and isolated genomes of *Akkermansia*

In this study, we obtained 1,119 *Akkermansia* MAGs conforming to the “near-complete” standard (completeness >90% and contamination <5%, **Supplement Table ST1**) from Pasolli *et. al*. study [20], aiming to decipher the population structure and geographical distribution of *Akkermansia*. Despite the metagenomic samples were collected extensively from multiple human body sites, we found that all *Akkermansia* MAGs were detected from the human fecal samples (**Supplement Table ST1**). This finding was in agree with the previous studies showing that the gut, rather than other body sites, is a major habitat of *Akkermansia* [21]. Similar, only 6 non-*Akkermansia* Verrucomicrobia genomes were identified in a recent study reconstructing over 56,000 MAGs from global human oral metagenomes [22]; this result also showed a very low occurrence of *Akkermansia* in human oral cavity.

The average completeness and contamination rates of 1,119 *Akkermansia* MAGs were 96.3% and 0.4%, respectively. The genomic data revealed varying genomic size ranging from 2.17 to 3.30 Mbp (averaging 2.73 Mbp, **Figure 1A**). The MAGs represented five continents and 22 different countries. The majority of genomes (62.4%, 698/1,119) were from countries in the European, and the others were from Israel (n = 141), America (n=100), China (n = 97), Russia (n = 33), Kazakhstan (n = 33), Mongolia (n = 10), Fiji (n = 5), and Peru (n = 2) (**Figure 1B**). In view of the geographical and population spans and the integrity of 1,119 MAGs, we suggested that they effectively represented the characteristics of the human intestinal *Akkermansia* genus and can be used to answer fundamental questions regarding population structure and functional specificity of *Akkermansia*.

**Figure 1.**
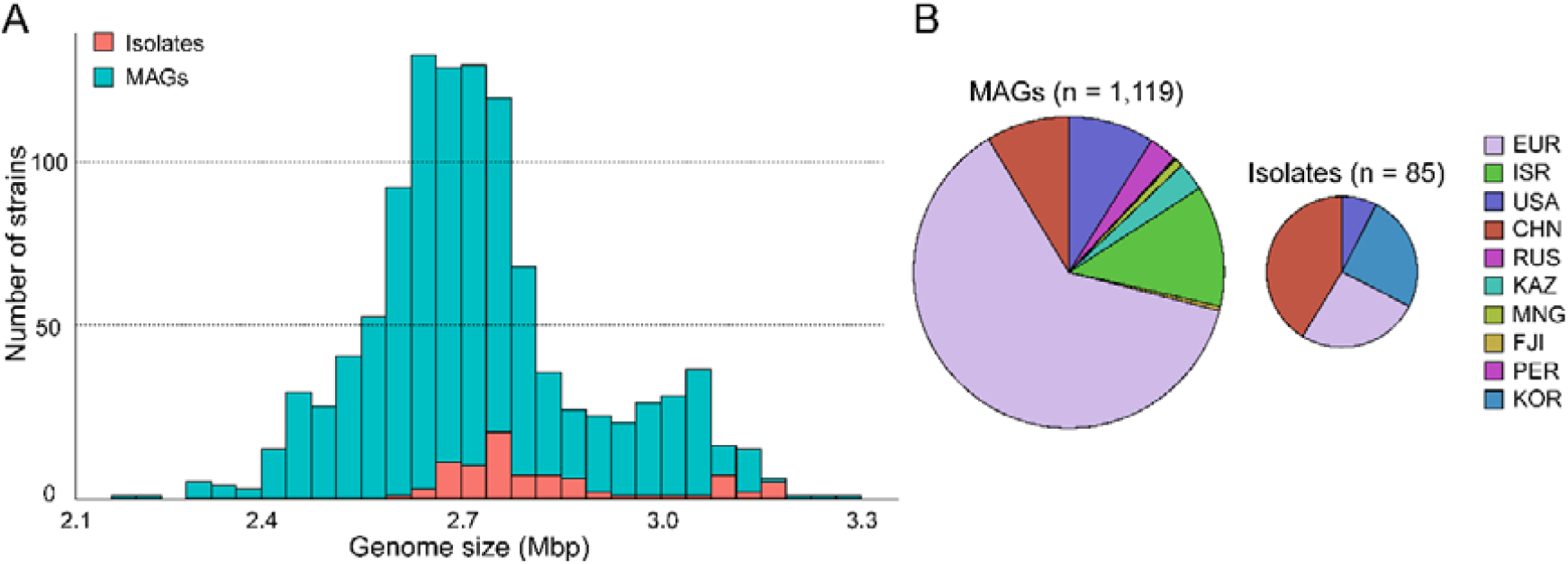
The genome size and country distribution of 1,204 strains of *Akkermansia*. **A**. Distribution of genome size of 1,119 MAGs and 85 isolated genomes. **B**. Country distribution of 1,119 MAGs and 85 MAGs isolated genomes.

To extent the genomic content of *Akkermansia*, we also analyzed 84 isolated genomes from National Center of Biotechnology Information (NCBI) database (up to Aug. 2019) and 1 newly sequenced *Akkermansia* strain (GP37, an *A. glycaniphila* strain isolated from human gut) (**Supplement Table ST2**). The distribution of genome sizes of the isolated genomes was consistent with that of the MAGs (**Figure 1B**). All of these strains were isolated from the feces, but their host were widely distributed, including human (n = 61), mice (n = 13), chimpanzee (n = 3), and other animals (n = 8). 96.5% (82/85) of the isolated genomes were *A. muciniphila*, and the remaining 3 were *A. glycaniphila*.

### Population structure of *Akkermansia*

The phylogenetic relationship of all 1,204 *Akkermansia* genomes was analyzed based on PhyloPhlAn, a method for improved phylogenetic and taxonomic placement of microbes [23]. We identified seven distinct phylogroups of *Akkermansia* (**Figure 2A**), including three *A. muciniphila* phylogroups (Amuc I, Amuc II, and Amuc III) previously reported by Guo *et. al*. [19] and a phylogroup of *A. glycaniphila*. The three *A. muciniphila* phylogroups accounted for 94% of all genomes, of which, 945 were Amuc I, 181 were Amuc II, and 6 were Amuc III.

**Figure 2.**
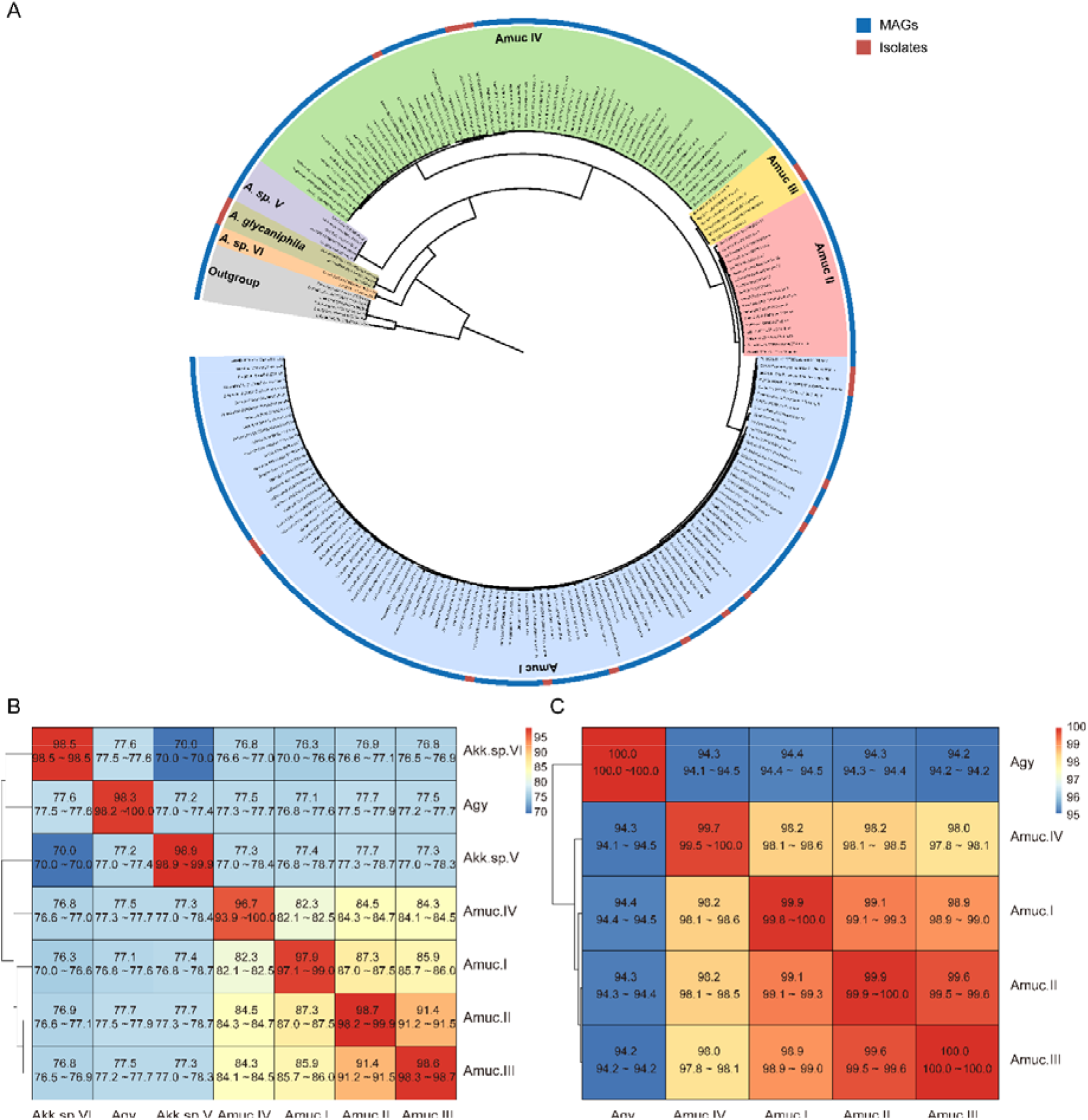
Phylogenetic analysis of *Akkermansia* genomes. **A**. Phylogenetic tree of 1,119 MAGs a85 isolated genomes. Filling colors in the phylogenetic tree represent different species or phylogroups. The outer circle represents the original of strains from MAGs and isolated genomes. For better visualization, only 10% strains of Amuc I and Amuc II were used, without changing the overall structure of the tree. **B-C**. Heatmaps show the pairwise ANI among 7 *Akkermansia* species and phylogroups **(B)** and the 16S sequence similarity among 5 *A. muciniphila* phylogroups and *A. glycaniphila* **(C)**.

The clustering result of average nucleotide identity (ANI) on whole genome data was identical with the phylogenetic analysis (**Figure S1**). The three known *A. muciniphila* phylogroups (Amuc I, II and III) were with average between-phylogroup ANIs ranging from 85% to 91% (**Figure 2B**) and average 16S rRNA genes similarity from 98.9% to 99.9% (**Figure 2C**). This finding suggested that these phylogroups were distinct subspecies, which was consistent the previous study [19]. In addition, a new phylogroup (containing 62 genomes) was with average ANI to these three *A. muciniphila* phylogroups of 82-84%, and average 16S rRNA genes similarity of 98.1-98.5%. This new phylogroup was thus defined as *A. muciniphila* subsp. IV (Amuc IV) followed the criterion of other *A. muciniphila* phylogroups [19]. Of *A. glycaniphila* and two ramianing new phylogroups, they showed a remarkable differed between-phylogroup ANI (<80%) and 16S rRNA gene similarity (<90%), suggesting that the were different species. These two new phylogroups were named as *Akkermanisia* sp. V (containing 5 genomes) and *Akkermanisia* sp. VI (containing 2 genomes).

There are significant differences in some genomic characteristics of the 7 *Akkermansia* phylogroups. Strains of *A. glycaniphila* and *Akkermansia* sp. V had the largest genome size (average 3.08 Mbp and 3.10 Mbp, respectively; **Figure 3**), and strains of *Akkermansia* sp. VI had the smallest genomes (average 2.39 Mbp). Of the *A. muciniphila* subspecies, the genome sizes of Amuc II were largest (average 2.99 Mbp), while Amuc I was smallest (average 2.66 Mbp). The distribution of number of proteins was consistent with that of genome sizes (**Figure S2**). The G+C content of *Akkermansia* sp. VI was extremely higher than others (average 64.8%; **Figure 3**), while that of *Akkermansia* sp. V was markedly low (average 52.0%). Amuc II and Amuc III had the higher GC content compared with other *A. muciniphila* phylogroups, and Amuc I had the lowest GC content. The representative genomes of Amuc I, Amuc II, Amuc III, and Amuc IV show differences in several different genomic regions (**Figure S3**). Diversified genomic characteristics of the *Akkermansia* phylogroups suggested different evolution history and functional habits of them.

**Figure 3.**
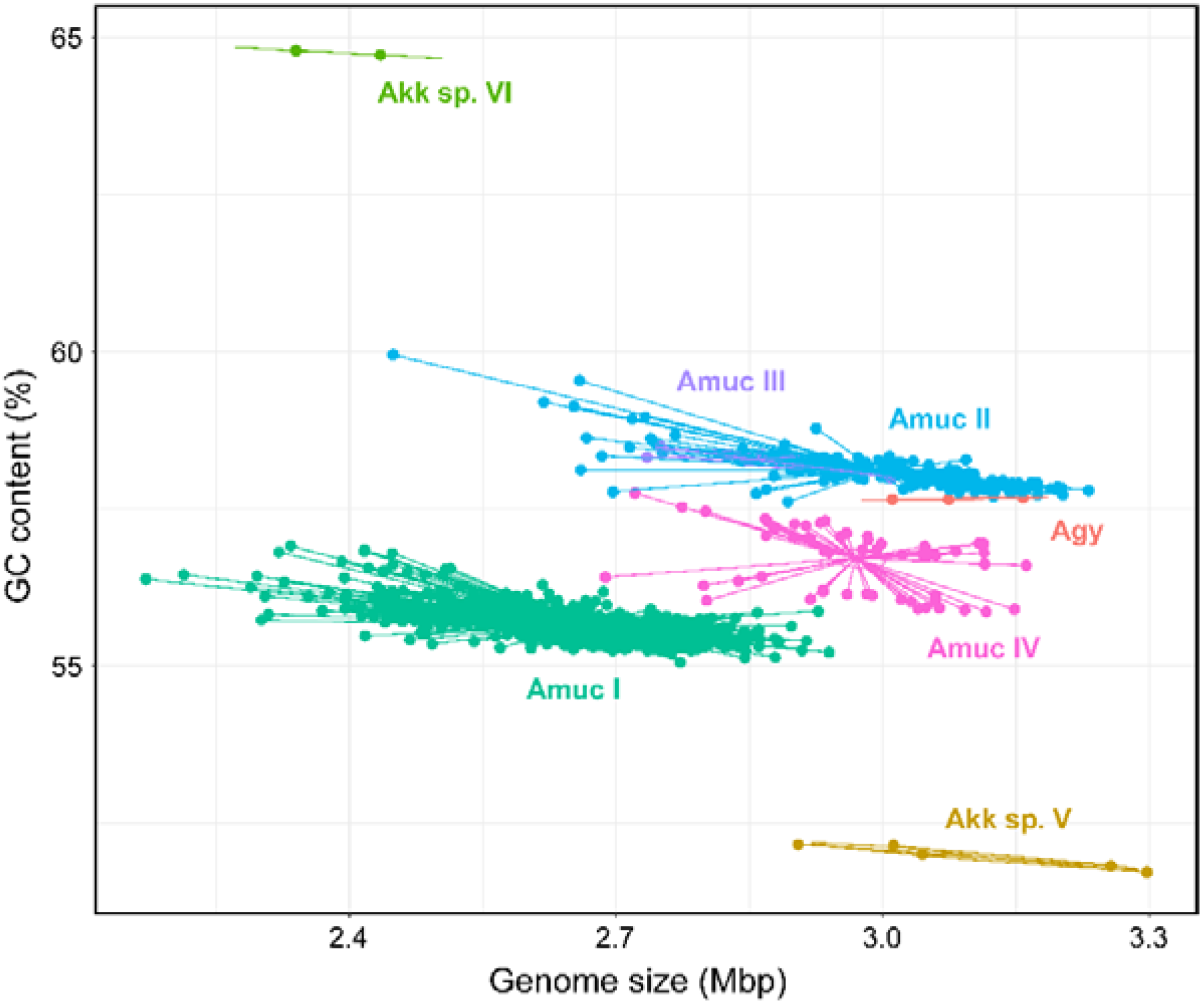
Scatter plot of the genome size and GC content of 7 *Akkermansia* species and phylogroups.

Our results expanded the known population structure of *Akkermansia* based on the most extensive collection of *Akkermansia* genomes at present. Belzer *et. al*. [6] divided the *Akkermansia* phylogenetic tree into five clades according to the full-length 16S rRNA sequences. In among, four clades of them contains human-associated sequences and one clade has highly diverse without human derived sequences. In this study, we included the *Akkermanisia* genomes from diverse transcontinental populations and found that the average similarity of 16S rRNA sequences between *A. muciniphila* and *A. glycaniphila* was 94.3% and among Amuc I-IV phylogroups was >98%. This result indicated a relatively conservative 16S rRNA sequence of *Akkermanisia* genomes in our study, suggesting more potential *Akkermanisia* species or phylogroups are still undiscovered, especially in non-human animals. A recent study [24] constructed the phylogenetic tree from 710 single-copy core genes shared by 23 *Akkermansia* genomes, and divided *A. muciniphila* into 4 sub-species. Their result was consisted with our findings showing that the genome of an *Akkermansia* strain (*Akkermansia* sp. KLE1797 from the NCBI databse) belonged to a distinct phylogroup (in our study, Amuc IV). Amuc IV was also found by Kirmiz *et. al*. based on 35 high-quality MAGs reconstructing from the feces of American children [25].

### Global distribution of *Akkermansia* phylogroups

The 1,204 *Akkermansia* genomes with wide distribution in 22 countries allowed us to investigate the biogeographical features of its phylogroups. We found that the two most dominant phylogroups, Amuc I and Amuc II, were extensively globally distributed (**Figure 4A-B**). Members of Amuc I and Amuc II were observed in trans-continental, trans-oceanic, cross-lifestyle populations, and even appeared across-host considering that all non-human *A. muciniphila* isolates were placed in these two phylogroups. Especially, Amuc II had a higher intra-phylogroup genetic diversity in western country compared to that of non-western populations (**Figure 4C**). On the other hand, the geographic bias of Amuc III and Amuc IV was more prominent (**Figure 4A; Figure S4**). Moreover, 83.3% (5/6) of Amuc III genomes were from the gut microbiotas of Chinese and only 1 genome was from European. Conversely, all 63 genomes of Amuc IV were from European, Israel and Peru, rather than from China or other countries. Moreover, the distributional modes of *Akkermansia* sp. V, *Akkermansia* sp. VI, and *A. glycaniphila* were still hard to estimate accurately because of the few number of genomes, however, all these species showed trans-continental distributed (**Figure S4**).

**Figure 4.**
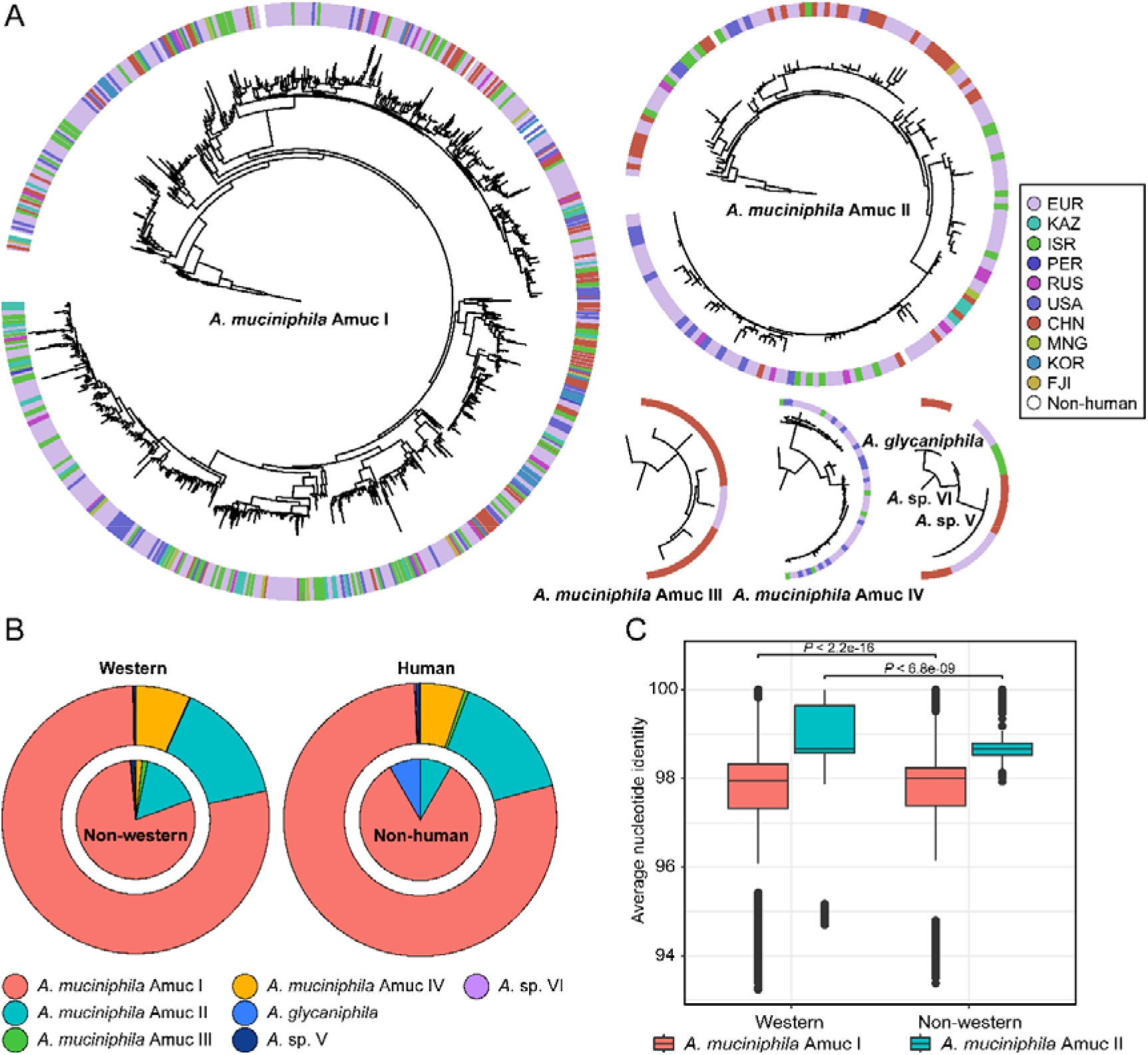
Geographic and host source of 7 *Akkermansia* species and phylogroups. **A**. Phylogenetic characteristics of *Akkermansia* species and phylogroups, with circles representing the national origin of each strain. **B**. The distribution of *Akkermansia* species and phylogroups in western and non-Western populations (left panel) and human and non-human host (right panel). **C**. Boxplot shows the intra-phylogroup ANI comparisons between western and non-Western strains.

Recently, the geographic deviation among different sub-species has been described in several gut bacteria such as *Prevotella copri* [26] and *Eubacterium rectale* [27]. We found that some phylogroups (Amuc I, Amuc II, and *Akkermansia* sp. V and VI) were global distributed with relatively low geographic deviation in *Akkermansia*, whereas Amuc III and IV were concentrated mainly in Chinese and non-Chinese populations, respectively. These findings suggested that not only geographical factor but also other unknown factors (maybe e.g. diet difference between populations [28], or movement of individuals) are the forces for speciation of the gut bacteria. Similarly, except for Amuc III, Amuc I, II, and IV, were observed in the gut microbiota of American children in Kirmiz *et. al*. study [25].

### Functional characteristics of *Akkermansia* phylogroups

It is potential that each *Akkermansia* phylogroup has a unique functional profile. To test this notion, we annotated all genomes using the Kyoto Encyclopedia of Genes and Genomes (KEGG) database [29] and identified a total of 1,568 KEGG orthologues (KOs) from them. PCoA analysis based on the KO profiles revealed a clear separation among three major phylogroups (Amuc I, Amuc II, and Amuc IV; *adonis* R^2^>0.25, *q*<0.001 in pairwise comparison) (**Figure 5A**), while the functions of Amuc III strains were relatively close to Amuc II but still significantly different (*adonis* R^2^=0.06, *q*=0.001). Likewise, the functions of *A. glycaniphila* and *Akkermansia* sp. V genomes were close to that of Amuc IV. We then compared the presence of KOs for three representative phylogroups (Amuc I, Amuc II, and Amuc IV) to identify the phylogroup-specific functions for them. In terms of pan-genome (KOs that occurred in at least one strain), the three phylogroups shared 1,026 functions, while 187, 64, and 40 functions were specific occurred in Amuc I, Amuc II, and Amuc III, respectively (**Figure 5B**). The Amuc I-specific functions involved to pathways of biosynthesis of secondary metabolites (20 KOs), ABC transporters (11 KOs), and two-component systems (8 KOs) and so on (**Supplement Table ST3**), while the Amuc II-specific functions referred to porphyrin and chlorophyll metabolism (10 KOs) and ABC transporters (7 KOs), glucose metabolism and others. In terms of core-genome (KOs that occurred in >90% strains), the three phylogroups shared 882 core functions, while 19, 29, and 30 functions were specific occurred in Amuc I, Amuc II, and Amuc III, respectively (**Figure 5C**). These phylogroup-specific functions were involved to multiple metabolism and transport pathways and potentially associated to the specific adaption mechanism for different *Akkermansia* phylogroups (**Supplement Table ST3**).

**Figure 5.**
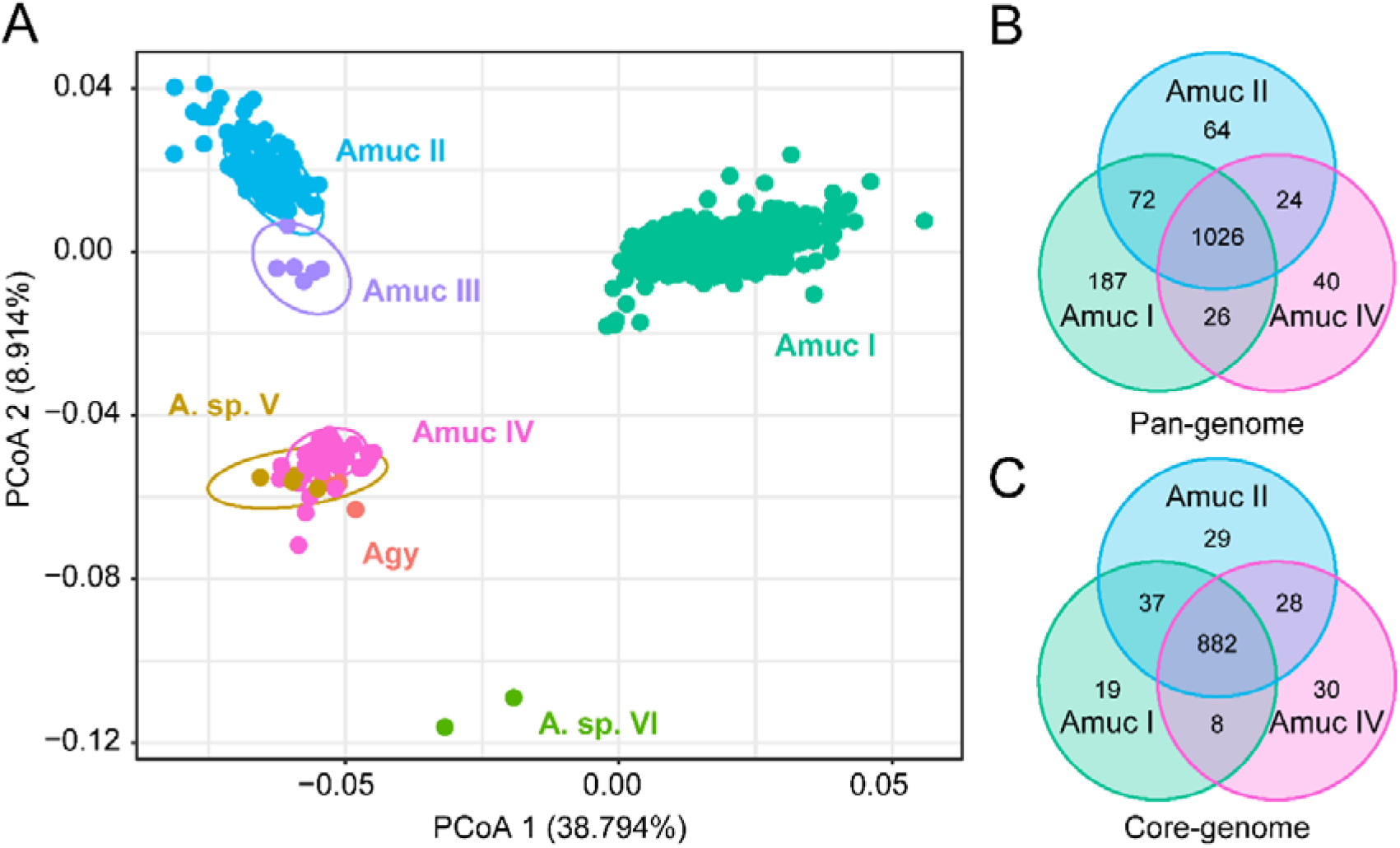
Comparison of KEGG functions among *Akkermansia* species and phylogroups. **A**. PCoA analysis on the KO profiles of 1,204 strains. **B-C**. Venn diagram shows the overlap of KOs in the pan-genome **(B)** and core-genome **(C)** of Amuc I, Amuc II, and Amuc IV.

In order to further investigate the ability of carbohydrates formation and decomposition of *Akkermansia* species, the genomes were screened for carbohydrate active enzymes (CAZymes) [30]. Noticeably, a remarkable separation was observed on the CAZyme profiles between members of Amuc I and Amuc II-III-IV (*adonis* R^2^=0.60, *q*<0.001; **Figure 6A**), while the strains of Amuc II, III, and IV were relatively closer. Compared to Amuc II-III-IV, strains of Amuc I had fewer total number of carbohydrate-active enzymes (*p*<0.001; **Figure 6B**) and especially glycosyltransferase (GT) proteins (*p*<0.001; **Figure 6C**). GT proteins are mostly related with protein glycosylation, cell wall polysaccharide synthesis or synthesizing exopolysaccharides in the context of biofilm formation [25]. This may represent the adaptability of the strains of Amuc II-III-IV phylogroups to the synthesis of exopolysaccharides or other structural carbohydrates. Moreover, 16 CAZymes were specifically encoded in the pan-genome of Amuc I members, the prominent of them were glycosyl transferase family 101 (GT101, occurred in 24.7% strains of Amuc I) and auxiliary activity family 7 (AA7, occurred in 5.6% strains); while 2 CAZymes belonged to carbohydrate esterase family 14 (CE14, occurred in 95.7% strains of Amuc II-III-IV) and glycoside hydrolase family 130 (GH130, occurred in 75.8% Amuc IV strains but none others) were specifically encoded in the genomes of Amuc II-III-IV (**Figure 6D; Supplement Table ST4**). GT101 is an β-glucosyltransferase that highly conserved in bacteria and involved to bacterial adhesion and biofilm formation [31]. CE14 involves in detachment of lignin from the polysaccharides that maybe lead to the function of cellulose hydrolysis in Amuc II-III-IV members [32]. Similarly, GH130 might potentially provide the ability of mannose hydrolysis for Amuc IV members [33] and this phenomenon may be correlated to the geographical distribution differences of the Amuc IV phylogroup. In brief, the difference of functional profiles between *Akkermansia* phylogroups may have relate to their ability for utilizing complex carbohydrates.

**Figure 6.**
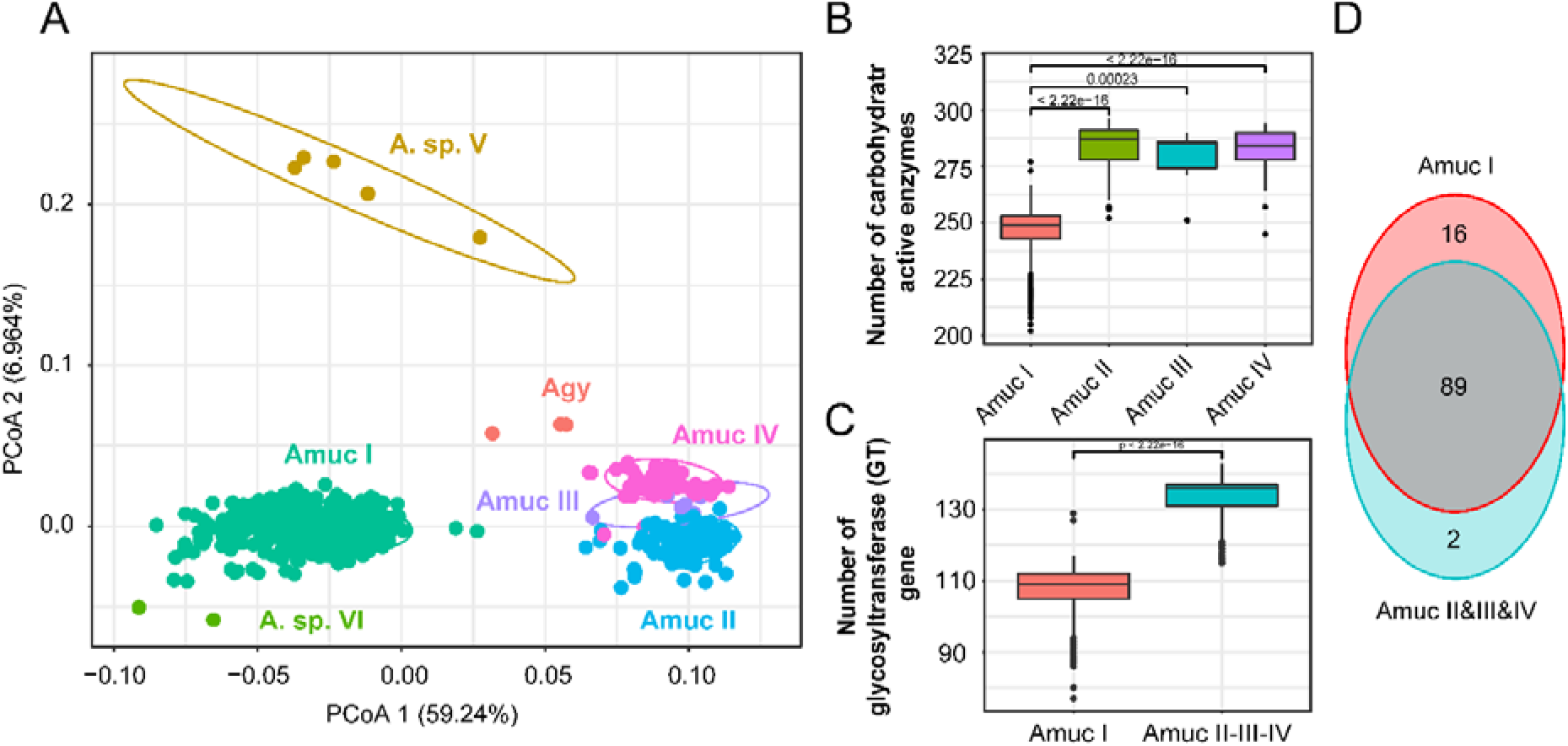
Comparison of CAZy functions among *Akkermansia* species and phylogroups. **A**. PCoA analysis on the CAZymes families of 1,204 strains. **B-C**. Boxplot showed the content comparison of CAZy enzyme-related genes annotated by the four phylogroups **(B)** and glycosyltransferase (GT) gene content between Amuc I and Amuc II-III-IV **(C)**, significance was calculated using the rank sum test. **D**. Venn diagram shows the overlap of CAZymes families between Amuc I and AmucII-III-IV.

## Conclusions

This study characterized the phylogeographic population structure and functional specificity of *Akkermansia* based on 1,119 near-complete MAGs and 85 isolated genomes. We placed the *Akkermansia* genomes into two previously isolated species (*A. muciniphila* and *A. glycaniphila*) and two candidate novel species (*Akkermansia* sp. V and *Akkermansia* sp. VI), and further divided the *A. muciniphila* members into three previously described phylogroups (Amuc I, II, and III) and a new phylogroup Amuc IV. These species and phylogroups revealed significant geographical distribution bias, especially Amuc III was presented in Chinese population and Amuc IV was mainly distributed in western populations. Functional analysis showed notable specificity in different *Akkermansia* species and phylogroups that involved to some metabolism and transport pathways and carbohydrate active enzymes. In conclusion, our results showed that the *Akkermansia* members in human gut had high genomic diversity and functional specificity and diverse geographical distribution characteristics.

## Methods

### Data source

1,119 MAGs of the *Akkermansia* were selected for analysis from the published dataset of Pasolli *et. al*. (http://segatalab.cibio.unitn.it/data/Pasolli_et_al.html) [20]. Each MAG met the quality standard of completeness more than 90% and contamination less than 5%, estimating by CheckM [34]. 84 sequenced *Akkermansia* strains were download from NCBI at Dec. 2019. An *A. glycaniphila* strain (GP37) was isolated from human feces and genomic sequenced using the Illumina HiSeq2500 instrument. Genomic assembly of *A. glycaniphila* GP37 was performed based on the previous pipeline [19]. The genome sequence of *A. glycaniphila* GP37 has been deposited in the NCBI BioProject PRJNA662466.

### Gene prediction and functional annotation

To unify the standards, all *Akkermansia* genomes were carried out genome content prediction using Prokka (v1.13.3) [35]. Prokka used a suite of tools to identify the coordinates of genomic features within sequences, including small rRNA (5S, 16S and 23S rRNA) using RNAmmer (v1.2) [36] and protein coding genes prediction using Prodigal [37] and the homologous proteins of known *Akkermansia* strains. Functional annotation of genes was performed via blasting against the KEGG (version Apr. 2019) [38] and CAZy databases (version Apr. 2019) using USEARCH [39] and DIAMOND [40], respectively, with parameters e-value <1e-10, identity >70%, and coverage percentage >70%.

### Bioinformatic analyses

A phylogenetic tree of the *Akkermansia* strains was produced from concatenated protein subsequences based on PhyloPhlAn [23] and visualized using iTol [41]. The outgroups of *Akkermansia* phylogenetic tree were 6 non-*Akkermansia* MAGs that belonged to Verrucomicrobia phylum. Pairwise ANI between two genomes was calculated using FastANI (v1.1) [42]. Statistical analyses were implemented at the R platform. Heatmap was performed using the heatmap.2 function, and principal coordinates analysis (PCoA) was performed using the cmdscale function (*vegan* package) and visualized using the *ggplot2* package [43]. The BRIG software was used to visualize of genome comparisons [44].

## Authors’ contributions

X.X.Z., Y.Z.P., S.H.L., Y.Z., and Q.B.L. conducted the study, performed the bioinformatic analyses, and wrote the manuscript. Y.C.W. edited the manuscript. All authors read and approved the final manuscript.

## Figure legend

**Figure S1.**
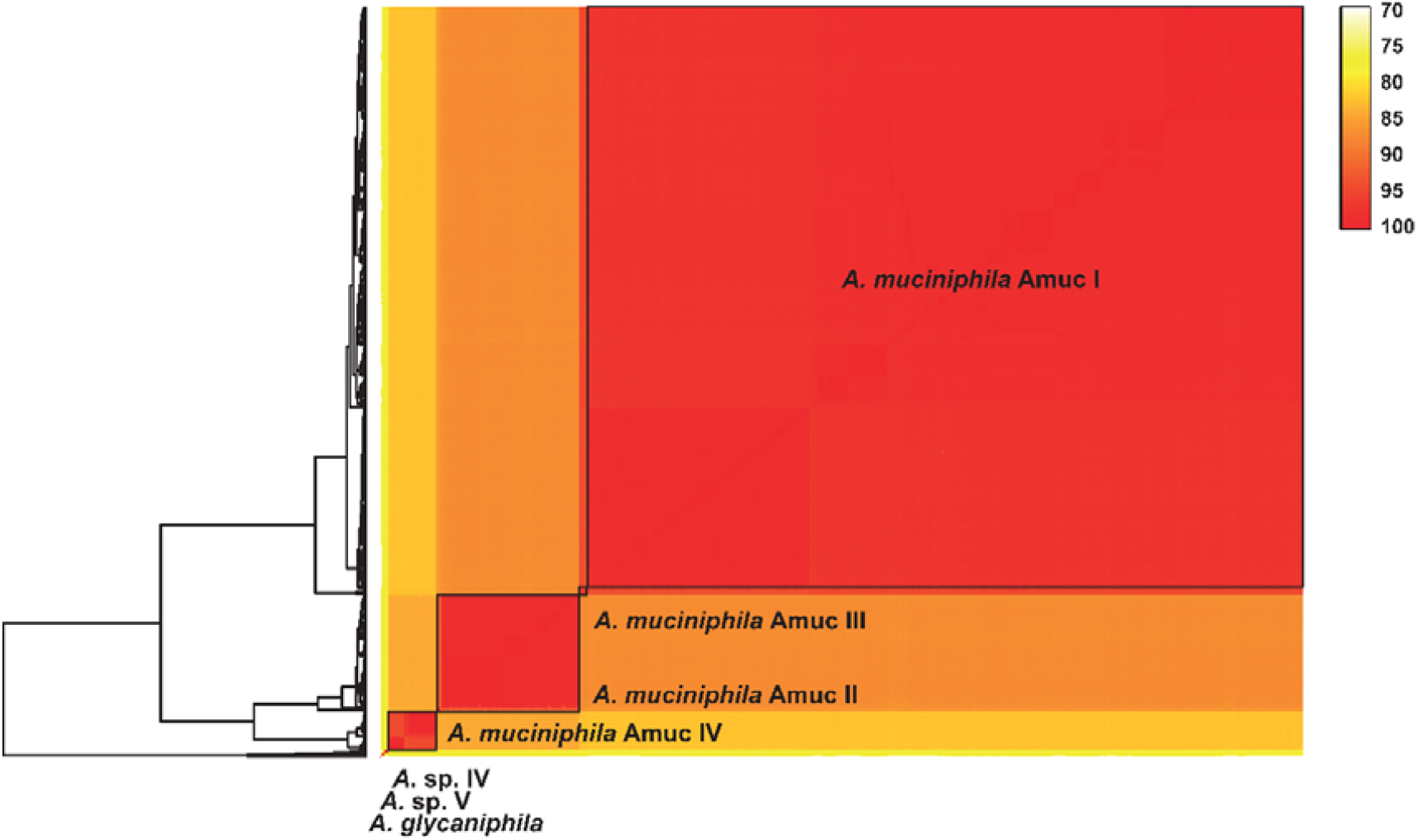
Heatmap shows the pairwise ANI values of 1,204 *Akkermansia* genomes.

**Figure S2.**
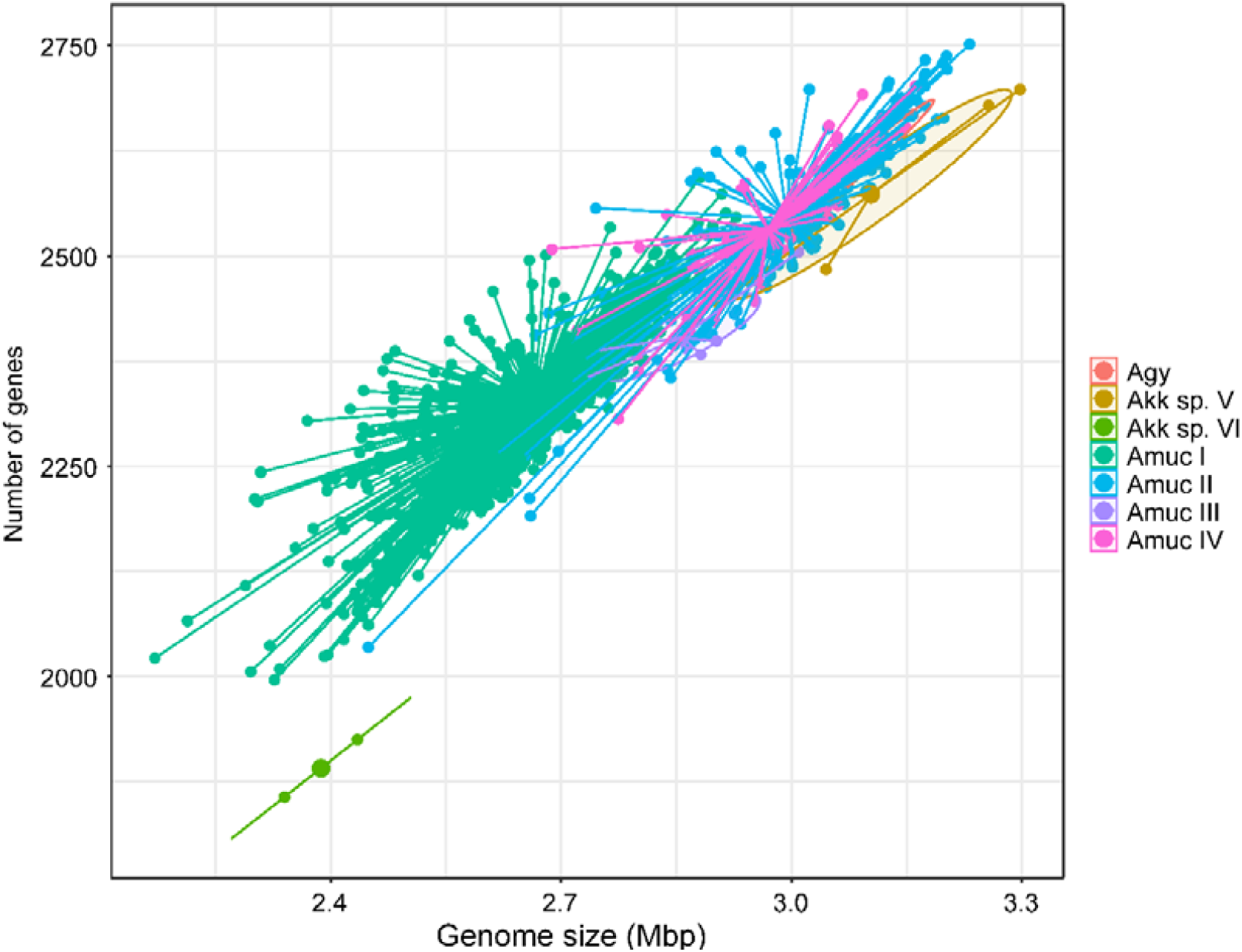
Scatter plot shows the relationship between genome size and gene content of 1,204 *Akkermansia* strain.

**Figure S3.**
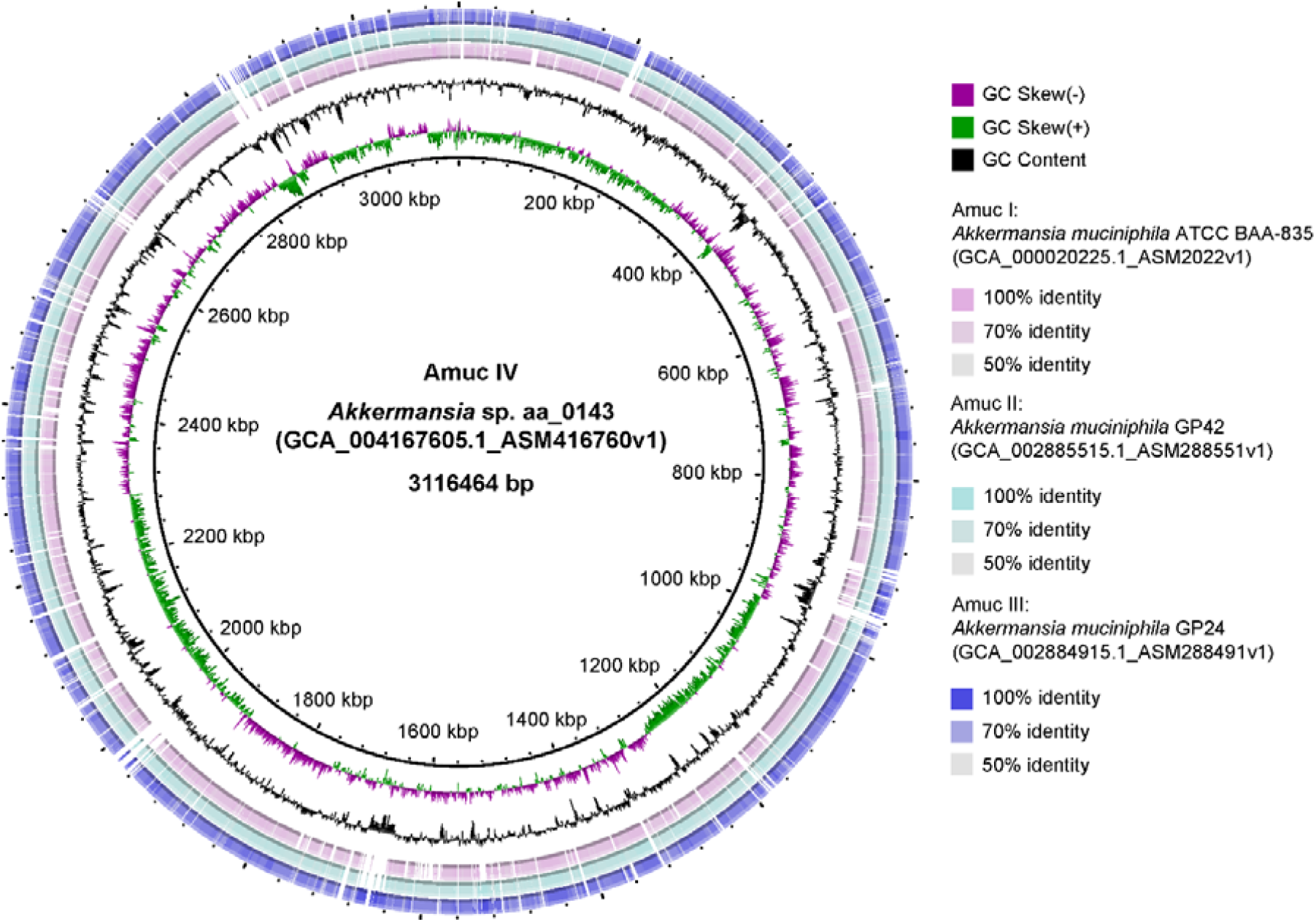
A genomic ring map of representative strain in Amuc IV phylogroup. The circle from inside to outside represents the Blast result of representative strain in Amuc I, Amuc II and Amuc III phylogroups with genome of Amuc IV phylogroup.

**Figure S4.**
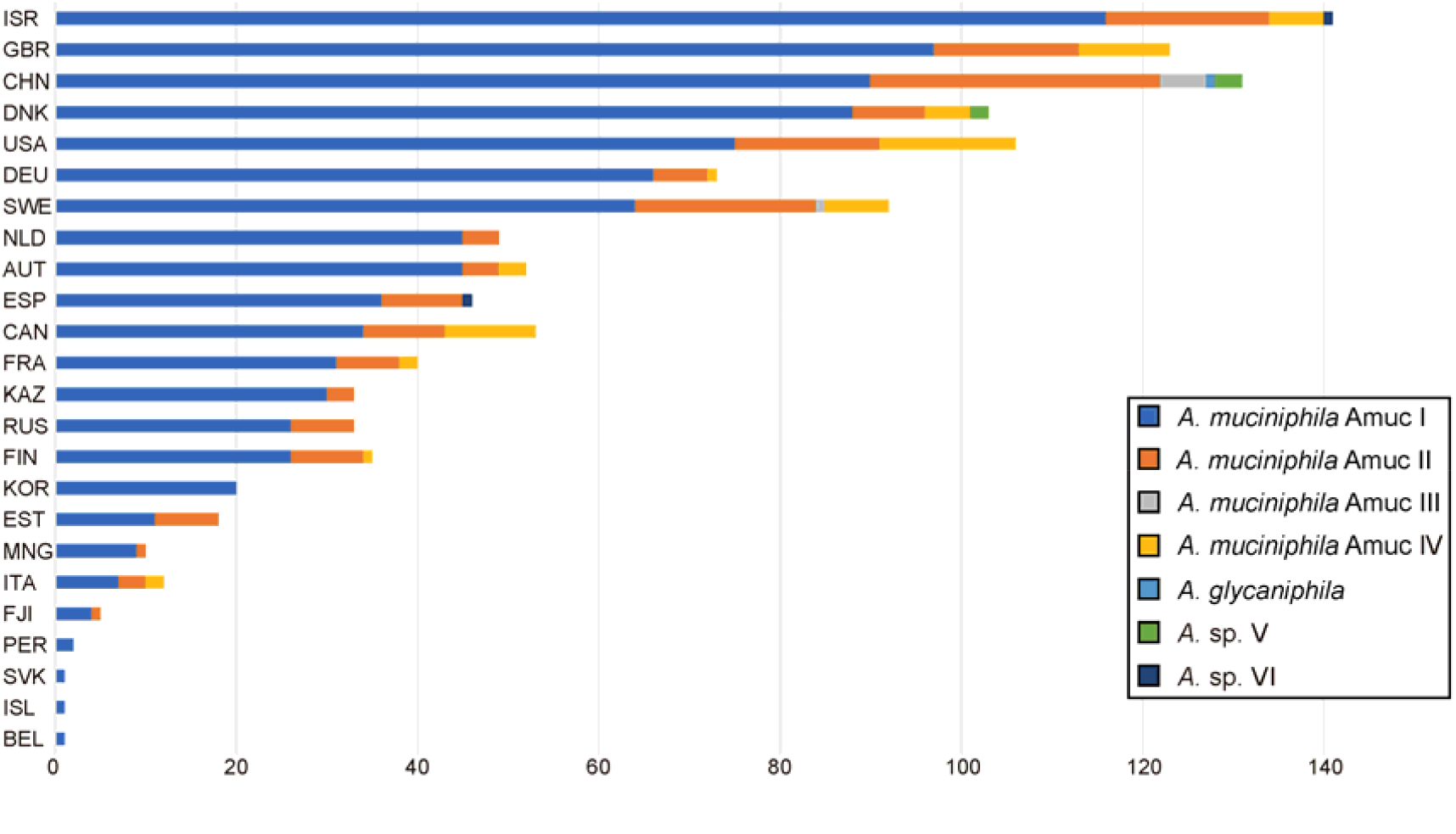
Country distribution of 7 *Akkermansia* species and phylogroups.

**Supplement Table ST1** The information of 1,119 MAGs of *Akkermansia*

**Supplement Table ST2** The information of 85 isolated genomes of *Akkermansia*

**Supplement Table ST3** The list of specific functions of *Akkermansia* phylogroups.

**Supplement Table ST4** CAZymes list of specifically encoded in the genome of *Akkermansia* phylogroups.

